# Biofilm and Cell Adhesion Strength on Dental Implant Surfaces via the Laser Spallation Technique

**DOI:** 10.1101/2019.12.11.873240

**Authors:** J. D. Boyd, C.S. Miller, M. E. Grady

## Abstract

**Objectives:** The aim of this study is to quantify the adhesion strength differential between an oral bacterial biofilm and an osteoblast-like cell monolayer to a dental implant-simulant surface and develop a metric that quantifies the biocompatible efficacy of implant surfaces.

**Methods:** High-amplitude short-duration stress waves generated by laser pulse absorption are used to spall bacteria and cells from titanium substrates. By carefully controlling laser fluence and calibration of laser fluence with applied stress, the adhesion difference between dental carry *Streptococcus mutans* biofilms and MG 63 osteoblast-like cell monolayers on smooth and rough titanium substrates is obtained. The Adhesion Index consists of a ratio of cell adhesion strength to biofilm adhesion strength obtaining a nondimensionalized parameter for biocompatibility assessments.

**Results:** Adhesion strength of 145±42 MPa is measured for MG 63 on smooth titanium, which increases to 288±24 MPa on roughened titanium. Adhesion strength for *S. mutans* on smooth titanium is 315±9 MPa and remained relatively constant at 332±9 MPa on roughened titanium. The Adhesion Index for smooth titanium is 0.46±0.12 which increased to 0.87±0.05 on roughened titanium.

**Significance:** The laser spallation technique provides a platform to examine the tradeoffs of adhesion modulators on both biofilm and cell adhesion. This tradeoff is characterized by the Adhesion Index, which is proposed to aid biocompatibility screening and could result in improved implantation outcomes. The Adhesion Index is implemented to determine surface factors that promote favorable adhesion of cells greater than biofilms. Here, an Adhesion Index >> 1 suggests favorable biocompatibility.

**Graphical Abstract:** 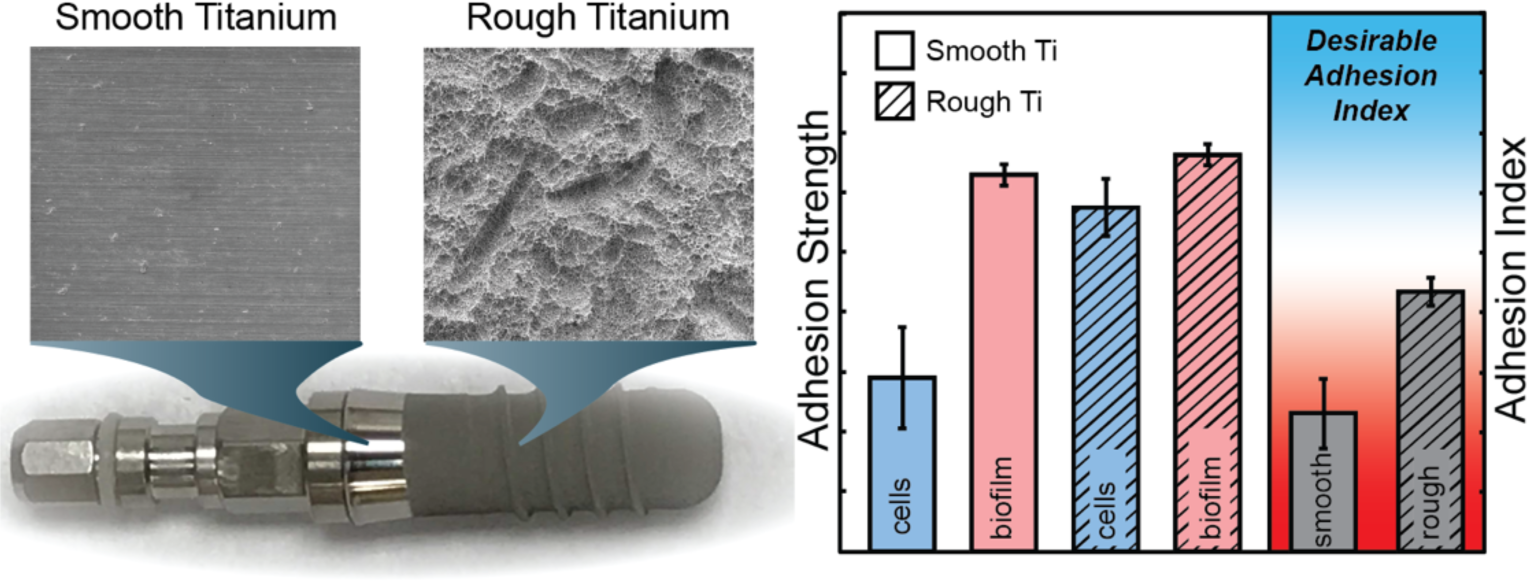

**Highlights:**

- Biofilm and cell monolayer adhesion are measured via the laser spallation technique
- Smooth and roughened dental implant-mimicking titanium surfaces are investigated
- Surface roughness increases cell adhesion but does not alter the adhesion of biofilms
- An Adhesion Index is developed to directly quantify the adhesive competition between bacteria and cells on an implant surface

## 1. Introduction

Biofilm formation is a significant problem in America, accounting for 17 million infections, causing 550,000 deaths, and costing the healthcare industry billions of dollars annually [1-4]. Bacterial biofilms are the leading source for chronic bacterial infections, due to the high antibiotic resistance of the biofilms [5, 6]. Complications from biofilm formation are prolific in implantology, accounting for half a million infections annually [7].

Biofilm formation is especially problematic with respect to dental implants. Dental implants are exposed to numerous oral bacteria, which can colonize the titanium surface leading to an infection called peri-implantitis. With infection rates as high as 28%, peri-implantitis is a serious problem in today’s dental community [8]. Peri-implantitis stems from the adhesion and development of a colonized bacterial biofilm onto the subgingival implant surface [9]. While there have been many advancements in the study of biomaterials, device related infections remain a critical problem. To prevent these bacterial biofilms from forming it is paramount to study and quantify the adhesion of bacteria onto various surfaces. Preventing the initial adhesion of pathogenic bacteria and biofilm formation would mark a significant step to deter bacterial infection of implants. Lack of available quantitative, high throughput, adhesion techniques hinders our progress toward optimal implant surface designs. Additionally, during implant design, biocompatibility assessments focus entirely on the implant-host response, omitting the impact of bacteria-implant-host response. An understanding of factors that contribute to strong biofilm adhesion at implant interfaces can guide the development of surfaces that prevent deleterious biofilms and promote osseointegration.

Unfortunately, there is still a large gap in knowledge of biofilm adhesion and the biocompatibility of implants, especially dental implants. Currently the most ubiquitous bacterial adhesion technique is quantitative polymerase chain reaction (qPCR) [10, 11]. While qPCR is excellent at quantifying the number of bacteria present on a surface, the technique lacks the ability to generate a quantified adhesion strength, which is related to force of removal. Critical force methods do exist such as atomic force microscopy (AFM), and jet impingement [12-16]. However, AFM is purely suited for pull-off forces for single bacteria, neglecting the effects of extracellular polymeric substances and macroscopic adhesion, and jet impingement tests over the entire biofilm, making it impossible to test a single biofilm more than once. The variety of testing methods also brings rise to a lack of consensus on the effects of surface roughness on bacterial adhesion. Some studies stating that it increases adhesion [17, 18], while other studies are unable to find a correlation [15, 19]. The lack of consensus on the effects on adhesion limits the development of optimized implanted devices. Another major problem with implant designs is there is no research which directly compares the adhesion strengths of bacteria and cells on similar surfaces. Current ISO-10993, the biological evaluation of medical devices, does not include the need for bacterial adhesion testing for implanted devices [20]. A direct comparison could aid in the bioassessments of implants as it will allow for an understanding of the tradeoffs for selecting different surface parameters. A bioadhesion assessment that compares the adhesion of both bacteria and host cells onto implant surfaces is needed.

In this work, the laser spallation technique is employed to measure the adhesion differential between bacterial biofilm and osteoblast-like cells on implant mimicking surfaces. The laser spallation technique is implemented to compare the effect of implant surface characteristics on bacterial biofilm, and cell monolayer adhesion in order to obtain quantitative adhesion measurements of each on rough and smooth titanium. The laser spallation technique receives macroscopic quantitative adhesion measurements and the localized stress wave loading allows for multiple testing on the same film. Titanium roughnesses are chosen to mimic those found on commercially available dental implants. The adhesion measurements for both host cells and deleterious bacteria can be compared directly to obtain an Adhesion Index. A single-species biofilm of *Streptococcus mutans* is chosen for the bacterial biofilm, and MG 63 osteosarcoma cells are chosen for the cell monolayer. *S. mutans*, a Gram-positive bacterium, is a major etiological agent of human dental caries that colonizes the oral cavity and forms bacterial biofilms [21]. Moreover, *S. mutans* has been shown to stimulate the growth and adhesion of deleterious bacteria and used in prior oral biofilm adhesion studies [15, 22, 23]. MG 63 osteosarcoma cells display numerous osteoblastic traits that are typical of immature osteoblasts that would adhere during osseous integration with the dental implant [24, 25]. Titanium is the current standard in the dental implant industry for many reasons such as its biocompatibility with bone and surrounding gum, high corrosion resistance, and its modulus of elasticity is comparable to that of bone [26]. Thus, commercially pure titanium is used to mimic the surface of a dental implant. Implant surfaces include roughened threading, to increase osseointigration, as well as unroughened surfaces. We selected both smooth titanium surfaces and roughened samples, with Ra 1.2 µm, which falls in the commercial standard of Ra 1-1.5 µm [19].

## 2. Materials and Methods

### 2.1. Substrate preparation

A complete substrate assembly is constructed to culture bacteria and cells while maintaining the integrity of the energy absorbing and confining layers needed for laser spallation [27]. Glass slides with one side coated with 100 nm of commercially pure titanium, 99.995% titanium, and the other side coated with 300 nm of aluminum are purchased from Deposition Research Laboratory Inc. (DRLI). The aluminum side of the sample is used as an absorbing layer for the Nd:YAG laser. A second set of slides are purchased from DRLI where the glass surface is sandblasted in order to achieve a uniform roughness of 1.2 µm, then coated in titanium. The surface roughness of the titanium is confirmed by a Zygo white light interferometer. Scanning electron microscope (SEM) images of the substrate assemblies are compared to the surfaces of a Straumann SLA dental implant in Fig. 1. Slides are cut into 1″x1″ squares and the aluminum layer is coated in a uniform thickness layer of sodium silicate (waterglass) (Fisher Chemical SS338-1) using a Specialty Coatings System G3P-8. These substrates are then adhered to the bottom of 35 mm Petri dishes with precut holes, using vulcanizing bioinert silicone (Dowsil 732 Multi-Purpose Sealant).

**Fig. 1.**
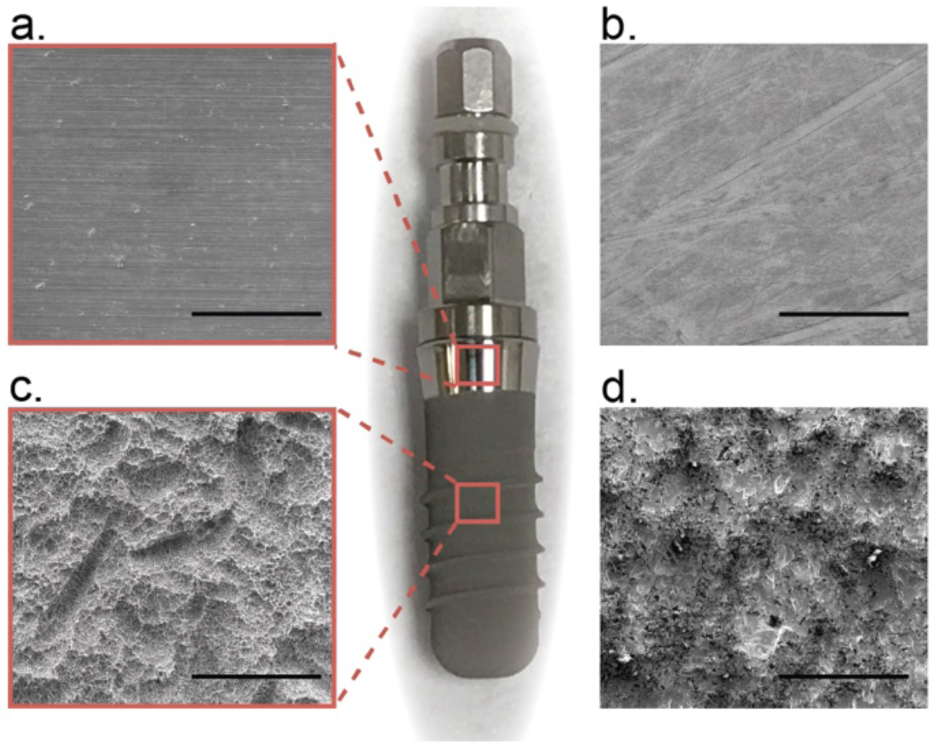
SEM images of Straumann dental implant surface (a,c) and dental implant-mimicking surfaces (b,d) for this study. Scale bars are 100 µm.

### 2.2. Cell and biofilm culture

*Streptococcus mutans* (Wild type Xc) [28] is cultured in Todd Hewitt Yeast broth (THY). *S. mutans* is cultured until an OD_600_ of 0.7 is obtained. The bacterial solution is added into the Petri-dish assemblies and diluted with a mixture of THY and 75 mM sucrose for a final OD_600_ of 0.175. Inoculated substrate assemblies are cultured at 37 °C with 5% CO_2_ and cultured for 24 hrs. Media is removed and the biofilms are gently rinsed with phosphate buffered saline (PBS) in order to remove any bacteria not colonized within the biofilm.

MG 63 (ATCC CRL-1427) is cultured inside a cell culture flask with Eagle’s Minimum Essential Medium (EMEM, ATCC 30-2003), 10% fetal bovine serum (FBS, ATCC 30-2020), 1% penicillin streptomycin solution (ATCC 30-2300) until confluent. The cells are then trypsinized and placed into an automatic cell counter. Cell concentrations of 120k are then placed inside the Petri-dish assemblies with more EMEM solution and incubated at 37°C with 5% CO_2_ for 48 hours, until confluent. After stress wave loading, biofilms and cells are dyed using Syto-9 (Thermo Fisher Scientific S34854) and Calcein AM (Thermo Fisher Scientific L3224), respectively, in order to determine viability of the surrounding cells. Films are then imaged using a Zeiss LSM 880 NLO upright confocal microscope. Z-stack images are then analyzed in biofilm thickness software, Imaris. The biofilms cultured on smooth titanium had an average thickness of 21.4±0.61 µm, and biofilms cultured on roughened substrates had an average thickness of 25.6±1.02 µm.

### 2.3. Laser spallation configuration and film loading

The laser spallation experimental setup used during biofilm and cell-substrate adhesion measurements is shown schematically in Fig. 2. An Nd:YAG laser pulse of 10 ns duration, wavelength of 1064 nm, with adjustable energy from 0 to 300 mJ, is used to obtain film spallation. A laser pulse is focused and reflected to impinge upon the backside of the substrate. Upon absorbing the laser energy, the sudden expansion of the absorbing layer generates a compressive stress wave that propagates towards the film on the front surface of the substrate. The wave then reflects at the thin film free surface resulting in a tensile load onto the biomaterial-titanium interface. Each substrate assembly is loaded at multiple locations by adjusting appropriate translation stages. The substrate assembly, depicted in Fig. 2b, and the experimental method of spallation testing are discussed in greater detail in Boyd *et al*.[27] and Kearns *et al*. [29].

**Fig. 2.**
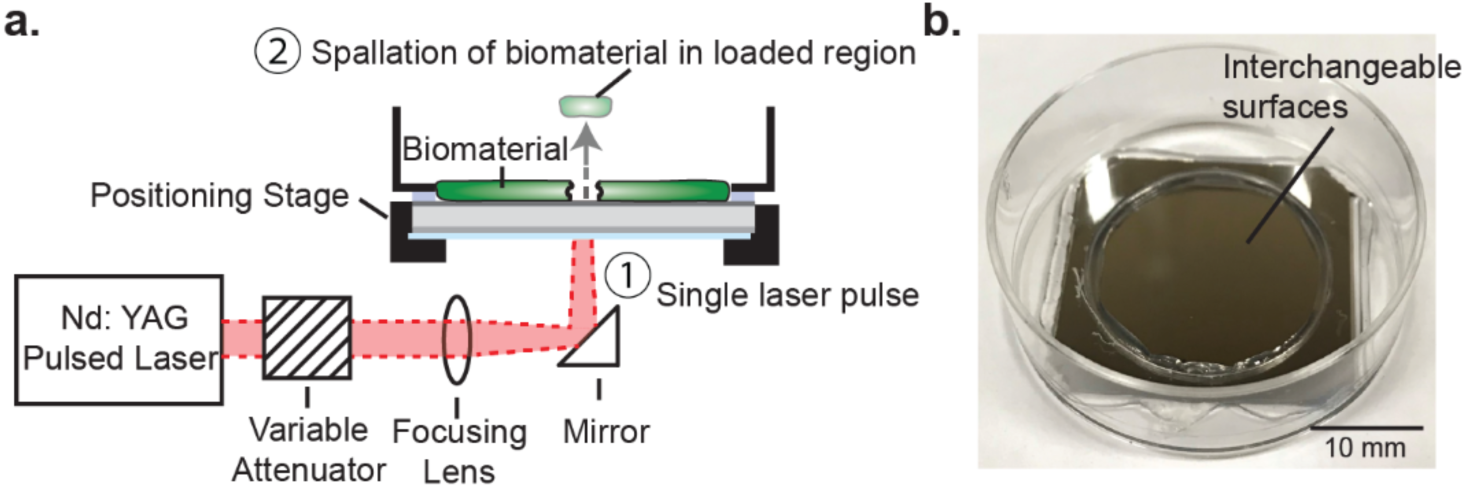
(a) Schematic of laser spallation setup used during experimentation where ➀ impingement of a single laser pulse ultimately initiates ➁ debonding of the biomaterial within the loaded region. (b) Substrate assembly before culture of test biomaterial.

During spallation testing both biofilm and cell monolayers are loaded over a range of fluences (7.93-79.4 mJ/mm^2^), which corresponds to 12-15 loading locations per test film. The experiment is repeated 12 times for each of the four conditions: *S. mutans* biofilm on smooth titanium, *S. mutans* biofilm on roughened titanium, MG 63 cells on smooth titanium, and MG 63 cells on roughened titanium. Overall, over 100 loaded regions are examined for each film, to determine fluence of failure. Failure is recorded when visible concentric ejection of the film at the loaded region is observed. The failure rate of each condition at each fluence is recorded, which is used to calculate the half-life and quantify adhesion strength.

### 2.4. Stress wave calibration

Stress wave calibrations are performed to convert laser energy to loading stress. Because biofilms and cells are nonreflective, *in situ* calibrations are precluded. Instead, calibration experiments are performed directly on unmodified substrate assemblies following previously described protocols [30, 31]. At each laser fluence, laser impingement and subsequent stress wave loading causes the surface of the substrate assembly to displace. These surface displacements are measured with a Michelson interferometer that includes a 532 nm continuous wave laser. Because the loading is rapid, over tens of nanoseconds, traditional displacement measurement devices are inadequate. A high rate oscilloscope (LeCroy WaveRunner 8404 M) records the temporal voltage trace from the Michelson interferometer via a silicon photodetector (Electo Optics ET 2030) shown by the equation,

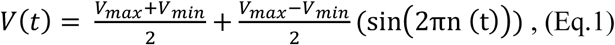

where V(t) is the voltage, V_max_, and V_min_, are the voltage maximum and minimum, and n(t) is the fringe number. From the voltage trace, the fringe number n(t) is unwrapped and converted to displacement, u(t), using (Eq. 2) and the wavelength of the interferometric laser, λ_0_ = 532 nm [32].

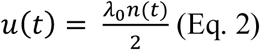

Fig. 3ab contain an example voltage trace for a single fluence and the corresponding displacement at that fluence alongside displacements for two other fluence values. In Fig.3b, lower fluence values results in less displacement when compared to the displacement of the higher fluence, which is expected. For fluence values of 39.71, 63.54, and 79.42 mJ/mm^2^, the resulting maximum displacement is 2, 4, and 5 µm. For a simple bi-material interface, the evolution of the substrate stress can easily be determined from the displacement history using the principles of one-dimensional wave mechanics [30]. Thus, using the displacement history, density of material ρ, and speed of sound through the material C_d_, the substrate stress profile, σ_sub_, is obtained by Eq.3.

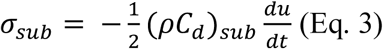

**Fig. 3.**
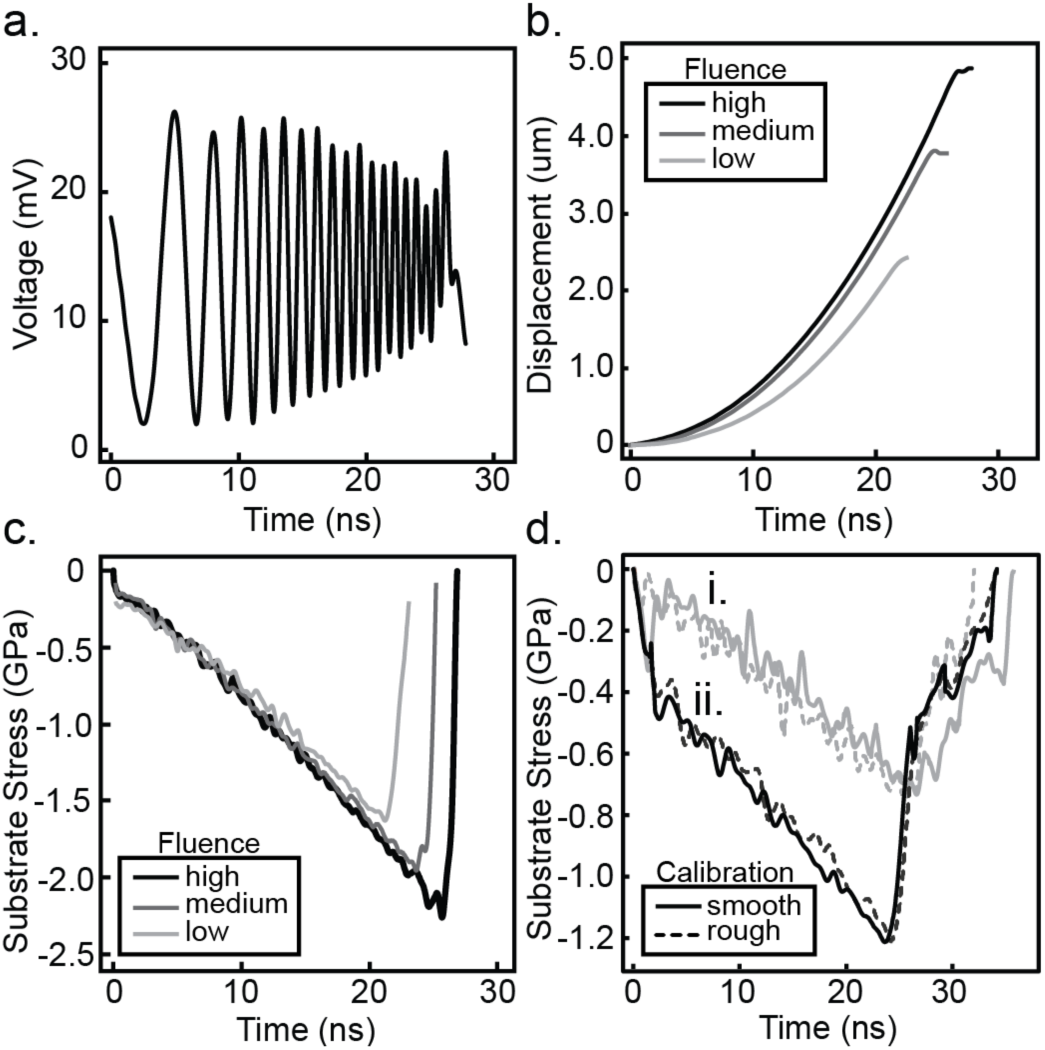
Raw data in (a) depicts a typical voltage curve recorded during calibration experiments for a high fluence, (b) depicts the displacement over time for a low, medium, and high fluence, and (c) depicts the associated substrate stress profiles calculated for the low, medium, and high fluence tested. Low, medium, and high fluences correspond to 31.8, 55.6, and 79.4 mJ/mm^2^, respectively. Raw data in (d) depicts the calibration between rough (dashed line) and smooth (solid line) substrates at a fluence of (i.) 55.6 mJ/mm^2^ in gray and (ii.) 79.4 mJ/mm^2^ in black.

Fig. 3c contains the substrate stress profiles obtained for the same three displacement profiles shown in Fig. 3b. An increase in laser fluence results in an increase in peak substrate stress. For fluence values of 39.71, 63.54, and 79.42 mJ/mm^2^, the resulting peak substrate stress is 1.5, 2, and 2.3 GPa. The slope of loading substrate stress profile, *i.e*., slope in the first 20 ns, for each fluence overlap each other, this result is expected since the slope is determined by the substrate material, glass in our case.

In order to perform calibration experiments on the roughened titanium, thin cover slips, 170 µm thickness (VWR micro cover glass No. 2), are adhered to the surface with Norland 60 Optical Adhesive and then coated in 150 nm of aluminum by Lesker physical vapor deposition (PVD). The same procedure is performed on smooth titanium substrates and the substrate stress profiles are compared in Fig. 3d. The shape of the measured stress pulses shows good agreement at each laser fluence. Peak substrate stress amplitude is equal at all fluences tested, varying by less than one standard deviation from the smooth titanium calibrations. Thus, the substrate stress profiles revealed that the rough surface had no measurable impact on wave propagation and thus smooth titanium is used for accurate stress wave calibration. By performing a set of calibration experiments, the peak substrate stress at each fluence tested is measured and shown in Fig. 4 as average and standard deviation of triplicate measurements.

**Fig. 4.**
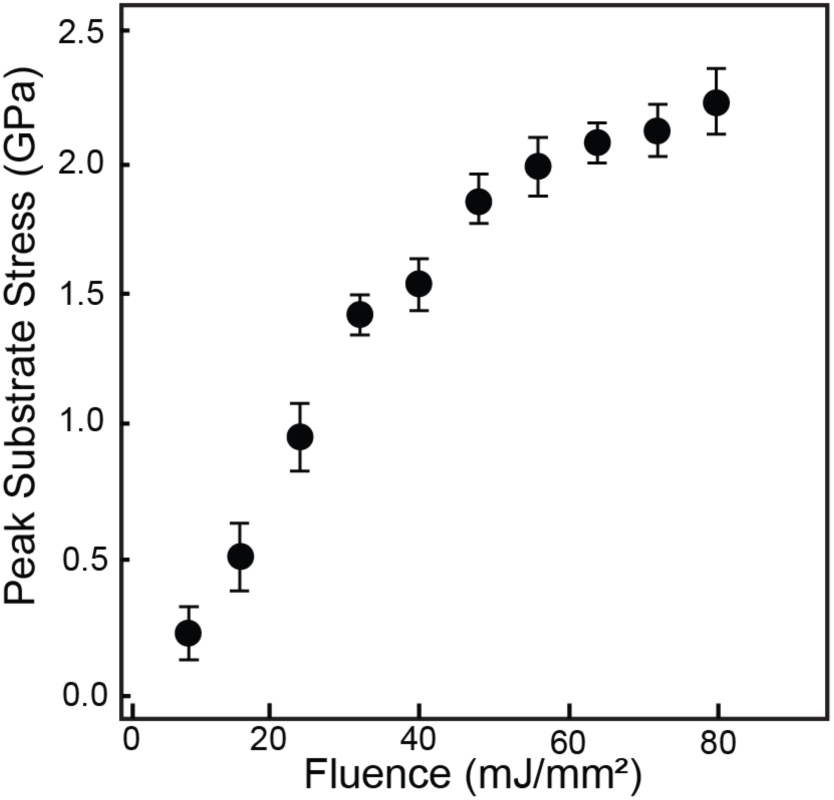
Average peak substrate stress measured at increasing laser fluence during spallation experiments. Error bars are one standard deviation.

Following the protocol developed by Kandula *et al*. [30] a modified equation for peak interface stress, σ_int,peak_, is derived using wave transmission and reflection coefficients,

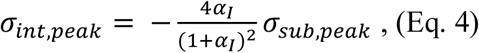

where α_I_ is equal to the ratio of the acoustic impedance, defined as the density times the dilatational wave speed, for the biofilm and titanium substrate, given as,

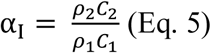

The density and dilatational wave speed of cells and bacteria for our calculations is assumed to be that of water, 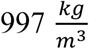 and 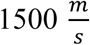, respectively, consistent with the works of other biomaterial researchers, including Paul *et al*. and Grover *et al*. [33, 34]. The density and dilatational wave speed for commercially pure titanium are 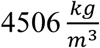 and 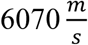, respectively. Through replacement of 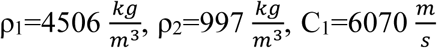, and 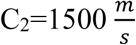 into Eq. 5 and substitution of α_I_ into Eq. 4, we obtain:

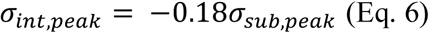

Thus, the peak interface stress is directly related to the peak substrate stress measured experimentally and determined by the loading laser fluence.

## 3. Results

### 3.1. Stress wave loading of biological films induces concentrated film ejection

*S. mutans* biofilms and MG 63 monolayers are loaded using the laser spallation technique. The loading results in high concentrated film ejection but leaves the surrounding cells adherent. The failure progression of each film tested is represented in Fig. 5. Fluorescent images were captured with a Nikon upright fluorescent microscope (Nikon Eclipse DS-Qi2) and a 10x objective. An upright microscope must be used because the titanium substrates are non-transparent. Images in Fig. 5 row 1 are from unloaded regions of each film. Fig. 5 row 2 and 3 include images of loading locations at a fluence of 39.7 mJ/mm^2^ and 79.4 mJ/mm^2^, respectively. Loading of MG 63 on smooth titanium at 39.7 mJ/mm^2^, row 2 column 1, results in film ejection while MG 63 on rough titanium at the same fluence, row 2 column 2, results in minimal film disturbance. Since the applied loading stress is the same at the same fluence, the difference in film failure is a direct result of the difference in adhesion strength. When comparing biofilm adhesion at the same fluence of 39.7 mJ/mm^2^, row 2 column 3-4, there is no film ejection. This difference indicates *S. mutans* biofilms have greater adhesion than MG 63 cell monolayers. Qualitatively, we found no noticeable effect on film loading for *S. mutans* biofilms on either smooth or roughened samples. At very high laser fluences, all films experience localized ejection (e.g., Fig. 5 row 3) while maintaining viability of the surrounding cells.

**Fig. 5.**
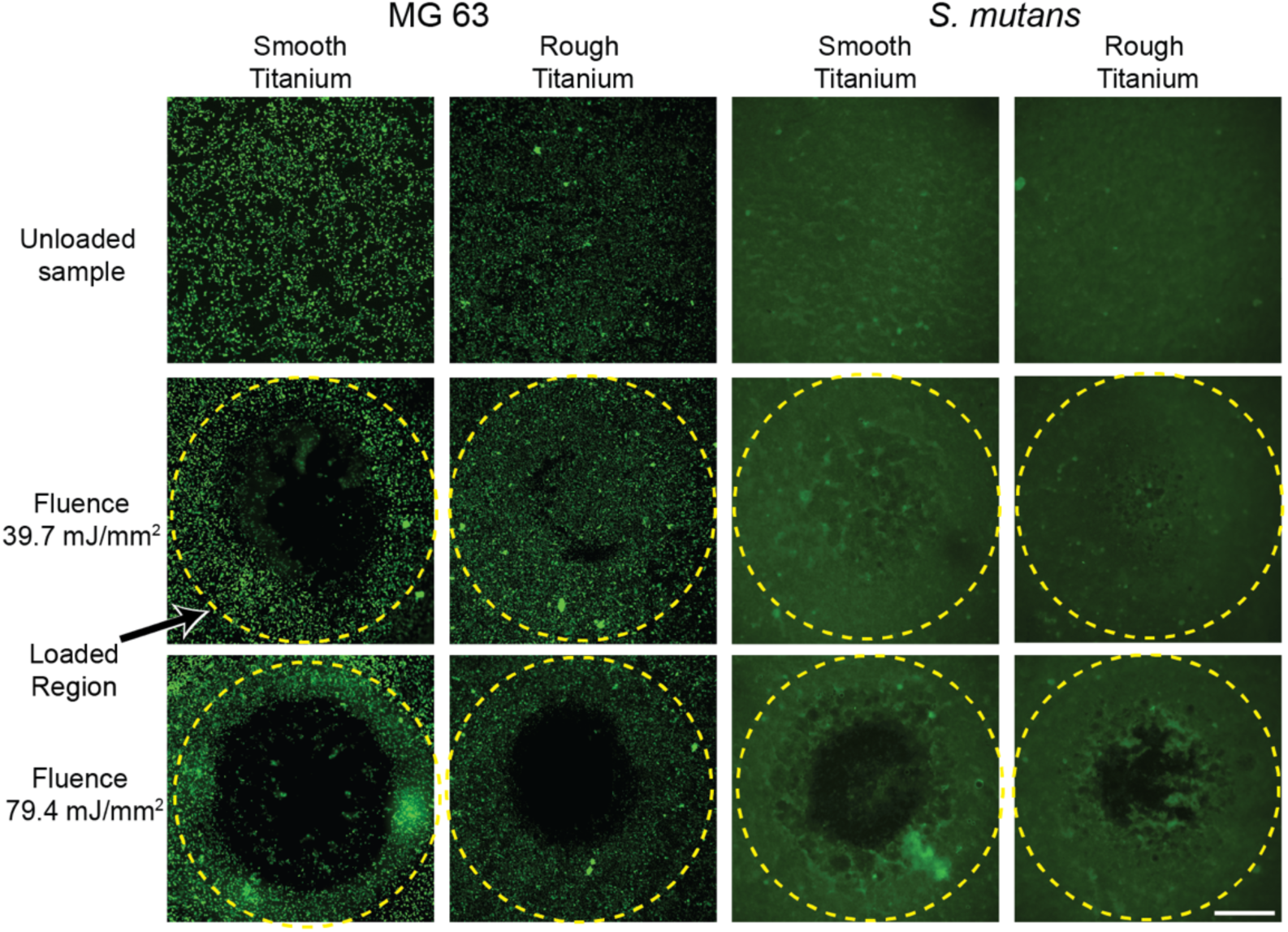
Fluorescence microscopy for MG 63 cell monolayers (first two columns on left) and *S. mutans* biofilms (last two columns) of an unloaded region (first row from top) and after loading at a fluence of 39.7 mJ/mm^2^ (second row) and a fluence of 79.4 mJ/mm^2^ (third row). Yellow dashed line indicates the loaded region. MG 63 cell monolayers and *S. mutans* biofilms are stained with Calcein AM, and Syto 9, respectively. Scale bar is 0.5 mm.

### 3.2. Adhesion strength determined by half-life failure statistics

Calibration experiments outlined in *Section 2.4* convert laser fluence values into interface stress for *S. mutans* and MG 63 films. Failure statistics recorded at each fluence across all replicates are combined (Fig. 6) in order to determine the adhesion strength of each film. In uniform homogenous films, the dichotomic presentation of film failure makes adhesion strength readily determined. However, the onset of film ejection, termed spallation, occurs over a range of fluence values instead of a single distinct fluence for biological films. For example, in Fig. 6a, at an interface stress of 272 MPa, 19% of biofilms on smooth titanium failed, while at an increased stress of 338 MPa, 89% of films failed. Biofilms grown on rough titanium saw a narrower onset of spallation and approached a more dichotomic relationship. The failure statistics, F(σ_int,peak_), are fit to a two parameter cumulative Weibull distribution function [35] (Eq. 7). Weibull analysis, common in macroscopic adhesion analyses [15, 36], calculates the half-life from a Weibull distribution, which is used as the adhesion strength, similar to the protocol developed by Grady *et al*. [37].

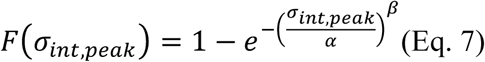

**Fig. 6.**
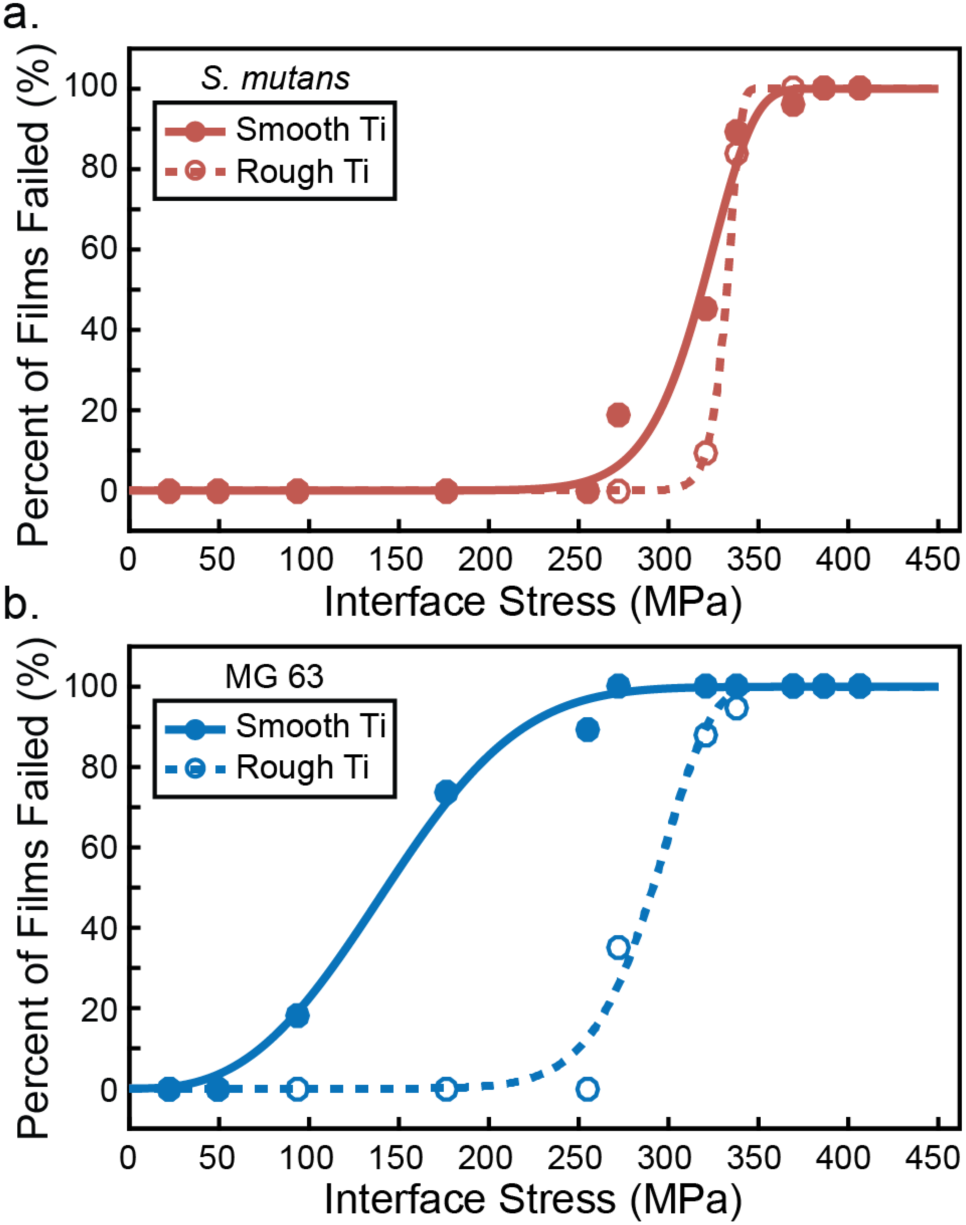
Failure statistics for (a) *S. mutans* biofilms on smooth titanium (solid red circles) and on rough titanium (open red circles) and (b) MG 63 on smooth titanium (solid blue circles) and on rough titanium (open blue circles) at increasing interface stress. Weibull models (smooth and dashed lines) are applied to interpolate the adhesion strength at a half-life of 50% failure.

The Weibull parameters, α and β, varied for each film condition and are included in Table. 1 as well as the root mean square (RMS) difference between the experimental data and the Weibull model. Weibull parameters are optimized to the lowest RMS value, the starred value in Table. 1 represents a local minimum which more closely approximated all of the experimental data points. The Weibull model is interpolated to obtain the adhesion strength at a half-life of 50% failure. The strength uncertainty is determined from the measured interface stress values.

**Table. 1.** Adhesion strength for each film condition, strength uncertainty, corresponding Weibull parameters, and root mean square (RMS) difference between Weibull model and experimental data.

| Film | Substrate | Adhesion Strength (MPa) | Strength Uncertainty (MPa) | $\alpha$ Parameter | $\beta$ Parameter | RMS |
| --- | --- | --- | --- | --- | --- | --- |
| *S. mutans* | Smooth | 315 | 9 | 326.5 | 15.02 | 0.0520 |
| *S. mutans* | Rough | 331 | 9 | 334.6 | 54.1 | 0.0006 |
| MG 63 | Smooth | 145 | 42 | 163.2 | 2.79 | 0.0281 |
| MG 63 | Rough | 288 | 24 | 300.1 | 12.31 | 0.0498* |

### 3.3. S. mutans biofilms exhibit higher interface adhesion strength than MG 63 osteoblast-like cells

Adhesion of *S. mutans* on smooth titanium is much greater than adhesion of MG 63 on smooth titanium. A qualitative comparison of images before and after loading for each film type from Fig. 5 reveals that the onset of spallation begins at lower stresses for MG 63 on smooth titanium compared to *S. mutans*. Film spallation has already occurred for MG 63 at 39.7 mJ/mm^2^, while no spallation is observed for *S. mutans* at the same fluence. The disparity in adhesion becomes more evident with our quantitative analysis of failure statistics and Weibull model in Fig. 6. The onset of spallation for MG 63 monolayers occurs at loading stresses greater than 50 MPa and saturates at 100% failure at loading stresses greater than 272 MPa. In stark contrast, a loading stress of 50 MPa does not induce separation of *S. mutans* biofilms from smooth titanium substrates. Failure for *S. mutans* does not occur until loading stresses reach 272 MPa and saturates at 100% failure at 387 MPa. The half-life value is obtained from the average value of the Weibull model for each cell and substrate combination. This half-life value is the adhesion strength and is plotted in Fig. 7. There is a two-fold difference in adhesion strength between MG 63 monolayers and *S. mutans* biofilms on smooth titanium. MG 63 has an adhesion strength of only 145±42 MPa, and *S. mutans* has an adhesion strength of 315±9 MPa.

**Fig. 7.**
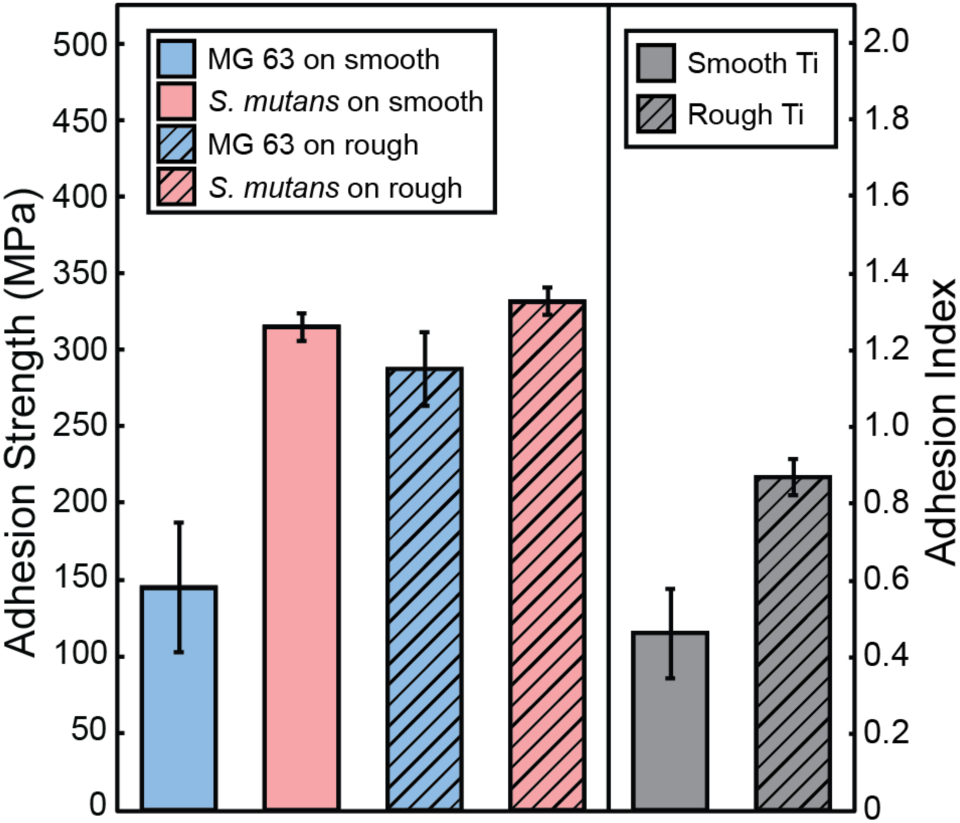
Adhesion strength for MG 63 (blue) and *S. mutans* (light red) biofilms on smooth (solid bars) and rough (hatched bars) surfaces. Surface roughness increases the adhesion for MG 63 with no effect on the adhesion strength of *S. mutans biofilms*. Adhesion Index, the ratio of MG 63 adhesion strength to *S. mutans* adhesion strength, is shown in grey for smooth and rough surfaces.

### 3.4. Titanium surface roughness increases adhesion strength of MG 63 monolayers, but not S. mutans biofilms

Similar to smooth titanium, the adhesion strength of *S. mutans* on roughened titanium is greater than MG 63 monolayers on roughened titanium, but MG 63 experiences a greater increase in adhesion compared to *S. mutans*. This result appears qualitatively through a comparison of loaded regions. For example, in Fig. 5 columns 2 and 4, images of MG 63 have very small regions where cells have ejected, while images of *S. mutans* show no film ejection. However, when comparing columns 2 and 4 with the images taken on smooth titanium, columns 1 and 3, a greater difference in spallation regions is observed for MG 63 monolayers. Additionally, when examining the failure statistics, the onset of failure for MG 63 drastically increases to 272 MPa (Fig. 6) on rough surfaces from 93.6 MPa, while the onset of failure *S. mutans* only increases to 320 MPa from 272 MPa. The adhesion strength for MG 63 increases to 288±24 MPa, and *S. mutans* adhesion strength increases to 331±9 MPa. While the adhesion strength obtained from the Weibull model, and displayed in Fig. 7, demonstrates that surface roughness increases adhesion for both films, the increase observed for MG 63 cell monolayer adhesion id drastically higher than the increase observed for *S. mutans* biofilm adhesion. These changes in adhesion strength correspond to a 98% increase in adhesion strength of MG 63 monolayers and only a 5% increase for *S. mutans* biofilms when smooth titanium is replaced by rough titanium. Therefore, surface roughness plays a bigger role in the adhesion of MG 63 cell monolayers when compared to *S. mutans* biofilms.

### 3.5. Surface roughness increases the Adhesion Index of titanium

In *Section 3.4*, we describe our finding that surface roughness affects adhesion of cell monolayers more than the adhesion of biofilms. To quantify the trade-off between increases in adhesion strength of cells and biofilms due to substrate modifications such as surface roughness, we developed the Adhesion Index. The ratio of the adhesion strength of cells (σ_cell_) to the adhesion strength of biofilms (σ_biofilm_) is the unitless Adhesion Index that describes which surfaces promote the adhesion of cells verses the adhesion of deleterious bacterial biofilms (Eq. 8).

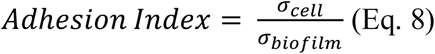

The adhesion strength of both films can be combined into the Adhesion Index, with the values for our experiments shown in Fig 7. When bacteria and cells are cultured onto smooth substrates the Adhesion Index is measured at 0.46±0.12, but when they are cultured onto roughened surfaces the Adhesion Index increases to 0.87±0.05. It is apparent by examining the Adhesion Index that roughening the titanium surfaces has a greater impact on cell adhesion than biofilm adhesion. Since neither smooth nor rough titanium is able to achieve an Adhesion Index greater than 1, there is room for improvement in implant designs to bridge this adhesion gap.

## 4. Discussion

In this work, high-amplitude short-duration stress waves generated by laser pulse absorption are used to spall bacteria and cells from titanium substrates. By carefully controlling the Nd:YAG laser energy fluence, the adhesion difference between *S. mutans* biofilms and MG 63 cell monolayers on both smooth and rough titanium substrates is obtained. The critical fluence value is converted into interface stress through established methods [30]. Our prior laser spallation adhesion studies of *S. mutans* biofilms demonstrated an increase in fluence of failure with increasing sucrose concentration [27], which was the first biofilm adhesion study by laser spallation. Other laser spallation experts have addressed mammalian cell adhesion. Hu *et al*. examined the adhesion of H19-7 neuron cells on silicon wafers [38], and Hagerman *et al*. examined MC3T3 fibroblast cells plated on untreated bacteria culture polystyrene [39]. The previous biofilm and neuron cell studies focused exclusively on fluence of failure, the measured critical fluence values in the range of 60 to 90 mJ/mm^2^ were measured for neurons on silicon, and a range of 20 to 50 mJ/mm^2^ was measured for biofilms on titanium. Hagerman *et al*. converted laser energy to interface stress and obtained adhesion values of 35 MPa for fibroblasts on polystyrene. Our test fluence values of 7.94-79.42 mJ/mm^2^ are consistent with other biological laser spallation results. We present the first study to directly compare adhesion measurements from laser spallation experiments of both bacteria and cells on the same surface.

The laser spallation technique has unique advantages for studying the macroscopic adhesion of biofilms due to its non-contact localized high strain rate force applied to cause film ejection. Despite such convincing evidence for the need for a macroscale critical force technique, there are limitations to this technique. For example, this technique is often interpreted as a way to clinically remove biofilms. That is not the goal of this work, though Meire *et al*. has written about the pros/cons regarding clinical adaptation [40]. Here we use the technique to measure adhesion strength to directly compare biofilm and cell adhesion. One limitation of this study is that it ignores the effects of co-culturing bacteria and cells, which could result in different adhesion results, and instead focuses on ideal adhesion for both cells and bacteria. Examining the ideal adhesion for both films something about the ratio of adhesion strengths through the Adhesion Index facilitates screening of surface tailoring, which includes blah blah and surface roughness.

Several studies have examined the effect of roughened titanium on adhesion of MG 63 cells [24, 41, 42]. By the 1980s, threads on dental implants were roughened because of improved osseointegration [43]. The adhesion studies that examine the effects of titanium surface roughness on cell adhesion have used counting methods to quantify adhesion and indicate that increasing roughness causes an increase in cell attachments, which correlates to increased osseointegration. Numerous studies have shown that implant surface roughness results in greater bone-to implant contact and higher resistance to torque removal [44-47]. However, there is no general consensus on the effect of surface roughness on adhesion of bacteria. For example, Aykent *et al*. [17] and Duarte *et al*. [18], who studied *Streptococcus sanguinis* and *S. mutans* report that increasing surface roughness results in increased adhesion. While Mei *et al*. [15], who studied *S. sanguinis*, indicate that surface roughness has no impact on bacterial adhesion. Similar roughness ranges of 1-2 µm were investigated in these experiments. Most of these studies that examine surface roughness effects rely on counting methods, with a few using AFM to measure pull off force for single bacteria. Shear flow studies have been conducted in the past to quantify the macroscopic adhesion of biofilms, but the increased time scale, several minutes, allows biofilms to adapt to surrounding forces [48]. Studies that only determine planktonic bacterial adhesion preclude macroscopic contributions of the EPS in biofilms towards adhesion. We believe the discrepancy within biomaterial adhesion studies of surface roughness is the result of at least three factors: 1) the use of a non-critical force adhesion measurement technique such as counting, 2) use of a micro or nanoscale adhesion technique to describe macroscale adhesive behavior or 3) the assumption that bacterial adhesion is the same as biofilm adhesion, which omits the contribution of biofilm EPS towards adhesion.

A major gap in the field of bioassessments is inability to directly compare bacterial and cell adhesion caused by the rampant use of non-quantitative adhesion techniques. Often, different techniques are employed to quantify both bacterial and cell adhesion. For example, Hauser-Gerspach *et al*. conducted adhesion studies using *S. Sanguinis* and *Porphyromonas Gingivalis*, alongside MG 63, however bacteria adhesion was quantified by enumerating colony forming units, and MG 63 adhesion was quantified by percent of surface covered by cells as well as nuclei counting [49, 50]. The lack of comparable techniques means that bacterial and cell adhesion are often studied separately and thus the effects of surface characteristics are not compared. This direct comparison is important as it will impact the efficacy of implantation. If a surface modification is shown to increase cell adherence and equally increase bacterial adhesion the risk of infection could increase. If bacterial adhesion occurs before tissue regeneration takes place, host defenses often cannot prevent surface colonization and formation of a protective biofilm layer [51, 52]. Therefore, it is critical to assess the predisposition of implant materials to strong biofilm adhesion. But how strong is too strong? This question is addressed by the introduction of the Adhesion Index.

The implementation of an Adhesion Index that directly compares the adhesion of host cells and deleterious bacteria, resulting in a nondimensional parameter will help weigh the effects of surface modifications on the relative adhesion strength between cells and biofilms. Fig. 8 illustrates the principles of the Adhesion Index. Values much less than 1 are undesirable as it indicates favoritism to bacterial biofilm adhesion. An Adhesion Index equal to one indicates that the adhesion strengths of cells and biofilms are equal. An Adhesion Index greater than 1 is desirable because that indicates the surface modification promotes cell adhesion over bacterial biofilm adhesion. Implementation of the Adhesion Index within our study indicates a more desirable Adhesion Index for roughened titanium over smooth titanium.

**Fig. 8.**
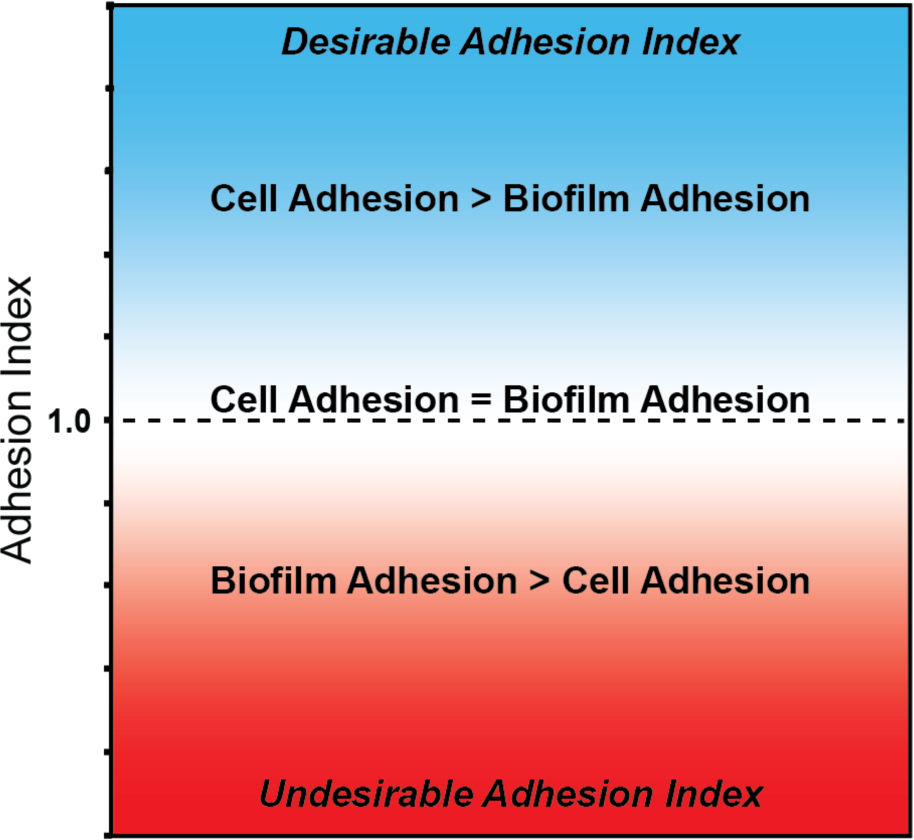
An ideal Adhesion Index demonstrates a much higher adhesion of mammalian cells than biofilms on a surface mathematically written as an Adhesion Index >> 1.

Current ISO-10993, the biological evaluation of medical devices, does not include the need for bacterial adhesion testing for implanted devices [20]. The Adhesion Index provides a target value to guide the design of new implant surfaces. We hypothesize that if measures of Adhesion Index are incorporated into biocompatibility assessment in addition to standard measures of cytotoxicity, genotoxicity, and other biological effects, we can improve the accuracy of preclinical assessment, which could lead to reduced propensity for biofilm forming infections on implant surfaces.

## 5. Conclusions

In this study, the laser spallation technique is implemented to measure the adhesion strength of *S. mutans* biofilms and MG 63 cell monolayers on titanium surfaces. The laser spallation technique is optimal as it introduces a localized non-contact force that captures the macroscopic adhesion effects for each film. The titanium surfaces selected simulate surfaces found on dental implants in order to investigate the effects of surface roughness on adhesion strength. Multiple adhesion tests are conducted for both *S. mutans* biofilms and MG 63 monolayers cultured onto smooth and roughened titanium substrates, and calibration experiments are performed to quantify adhesion strength. When increasing the titanium surface roughness, a significant increase in adhesion was measured for MG 63 monolayers, 145.3±42 MPa to 287.9±24 MPa, while a significant change in *S. mutans* biofilm adhesion was not observed, 315.3±9 MPa to 331.1±9 MPa. The adhesion values for MG 63 and *S. mutans* are directly compared to develop an Adhesion Index which quantifies the adhesive competition between the bacteria and cells on an implant surface. The Adhesion Index for smooth titanium is calculated as 0.46±0.12 and increases to 0.87±0.05 for roughened titanium. The nondimensional parameter, determined from the Adhesion Index, can help weigh the effects of surface modifications on the relative adhesion strength between cells and biofilms, and hopefully improve the efficacy of medical implant designs. The benefits of using our laser spallation technique setup is the ability to readily change surface and culture conditions, as well as bacteria and cell selection. The substrate assembly dishes can be swapped to examine a multitude of surfaces including other metals or even plastics and ceramics that might be used in other permanent (hip, knee) or temporary (catheter, tube) implants. Bacteria and culture conditions can also be modified to better match implant specific bacterial threats. Future work should expand the Adhesion Index to quantify the effect on adhesion for a variety of surfaces and using a multitude of bacterial and cell models.

## Research Data

The raw and processed data required to reproduce these findings are available to download from https://doi.org/10.18126/TW5W-XTWE [53] via the Materials Data Facility [54, 55].

## Acknowledgments

We acknowledge NIH COBRE Phase III pilot funding under number 5P30GM110788-04 to carry out these experiments. SEM images were taken within the Electron Microscopy Center at the University of Kentucky by staff associate Nicolas Briot. Aluminum sputter coating was performed at The Micro/Nano Technology Center at University of Louisville and overseen by Dr. Thomas Berfield. We thank the Center for Pharmaceutical Research and Innovation (CPRI) for use of bacterial culture equipment. CPRI is supported, in part, by the University of Kentucky College of Pharmacy and Center for Clinical and Translational Science (UL1TR001998). We thank Dr. Larissa Ponomareva for sharing her culture expertise.

